# ETV4 and ETV5 Orchestrate FGF-Mediated Lineage Specification and Epiblast Maturation during Early Mouse Development

**DOI:** 10.1101/2024.07.24.604964

**Authors:** Claire S. Simon, Vidur Garg, Ying-Yi Kuo, Kathy K. Niakan, Anna-Katerina Hadjantonakis

## Abstract

Cell fate decisions in early mammalian embryos are tightly regulated processes crucial for proper development. While FGF signaling plays key roles in early embryo patterning, its downstream effectors remain poorly understood. Our study demonstrates that the transcription factors *Etv4* and *Etv5* are critical mediators of FGF signaling in cell lineage specification and maturation in mouse embryos. We show that loss of *Etv5* compromises primitive endoderm formation at pre-implantation stages. Furthermore, *Etv4/5* deficiency delays naïve pluripotency exit and epiblast maturation, leading to elevated NANOG and reduced OTX2 expression within the blastocyst epiblast. As a consequence of delayed pluripotency progression, *Etv4/5* deficient embryos exhibit anterior visceral endoderm migration defects post-implantation, a process essential for coordinated embryonic patterning and gastrulation initiation. Our results demonstrate the successive roles of these FGF signaling effectors in early lineage specification and embryonic body plan establishment, providing new insights into the molecular control of mammalian development.

**Summary statement:** FGF signaling effectors ETV4/5 regulate lineage specification and embryo patterning in mice, affecting primitive endoderm formation and pluripotency exit.

## Introduction

FGF signalling is essential for mouse pre-implantation development where it acts in multiple successive steps. At the blastocyst stage, within the inner cell mass FGF drives the specification of primitive endoderm progenitors, an extra-embryonic endoderm lineage that later forms the yolk sac. Culture in exogenous FGF ligands directs all inner cell mass progenitors to differentiate to primitive endoderm (Nichols et al., 2009; Yamanaka et al., 2010). Conversly, inactivation of FGF signalling through genetic or biochemical means generates embryos with an inner cell mass containing all epiblast cells (Chazaud et al., 2006; Kang et al., 2017; Kang et al., 2013; Krawchuk et al., 2013; Molotkov et al., 2017; Nichols et al., 2009; Yamanaka et al., 2010). However, these epiblast cells have sustained elevated NANOG levels, indicating an inability to exit naïve pluripotency (Kang et al., 2017; Kang et al., 2013; Molotkov et al., 2017; Nichols et al., 2009). FGF signalling is well known to regulate naive pluripotency exit and later germ-layer differentiation *in vivo* (Lanner and Rossant, 2010; Nichols et al., 2009), and in mouse embryonic stem cells (Kunath et al., 2007; Ying et al., 2008). However, the direct targets and molecular mechanism of how FGF/ERK signalling executes cell fate decisions is poorly understood.

*Etv4* and *Etv5*, are ETS transcription factors, and downstream transcriptional activators of the FGF signaling pathway during embryonic development, with cooperative roles in the morphogenesis of the lung, limb bud and kidney (Herriges et al., 2015; Zhang et al., 2009). ETS transcription factor expression is induced by RTK/MAPK signalling, and post-translationally activated by MAPK phosphorylation (Charlot et al., 2010; Janknecht et al., 1996; O’Hagan et al., 1996). *Etv4* homozygous males are viable, but sterile (Laing et al., 2000), or have neuronal defects (Livet et al., 2002). Whereas, *Etv5* homozygous mutants die soon after birth (Zhang et al., 2009), or have mid-gestation (E8.5) lethality (Lu et al., 2009), the causes of which have not been established.

Single-cell transcriptomics of mouse (Ohnishi et al., 2014) and human (Blakeley et al., 2015) blastocysts revealed expression of *Etv4* (epiblast and primitive endoderm) and *Etv5* (inner cell mass progenitors and epiblast), which are downregulated in *Fgfr1* mutant mouse blastocysts (Kang et al., 2017), suggesting these ETVs may govern the transcriptional output downstream of FGF signaling in the inner cell mass. However, despite being implicated in regulation of the pluripotent state *in vitro* (Akagi et al., 2015; Kalkan et al., 2019; Zhang et al., 2018), the role of these transcription factors during epiblast development *in vivo* has not been assessed.

Here, we investigated the role of *Etv4* and *Etv5* as potential downstream effectors of FGF signalling in the establishment and maturation of the epiblast lineage. Analysing compound mutant mouse embryos, we find that loss of *Etv5* compromises the formation of the primitive endoderm at pre-implantation stages. While, at peri-implantation the loss of *Etv4/5* causes a delay in the progression of pluripotency and epiblast maturation, leading to developmental delay and AVE migration defects at early post-implantation stages. Together, our work sheds light on the successive roles of FGF signalling effectors in mediating inner cell mass fate decisions, and later epiblast maturation required for establishing the embryonic body plan at gastrulation.

## Results

### *Etv4* and *Etv5* expression during mouse embryonic development

Given the role of PEA3 family members in the regulation of the pluripotent state in stem cells, we hypothesised that PEA3 family member ETS transcription factors were involved in the establishment of the epiblast lineage. We first characterised where ETV transcripts were expressed during early mouse embryo development using our scRNA-seq dataset from mouse pre- to early post-implantation embryo development (Nowotschin et al., 2019). *Etv4* transcripts were expressed in late-blastocyst stage primitive endoderm (PrE) and epiblast (Epi), whereas, *Etv5* transcripts were expressed in early-blastocyst stage inner cell mass (ICM) progenitors, and later in epiblast cells (Fig. 1A).

**Figure 1:**
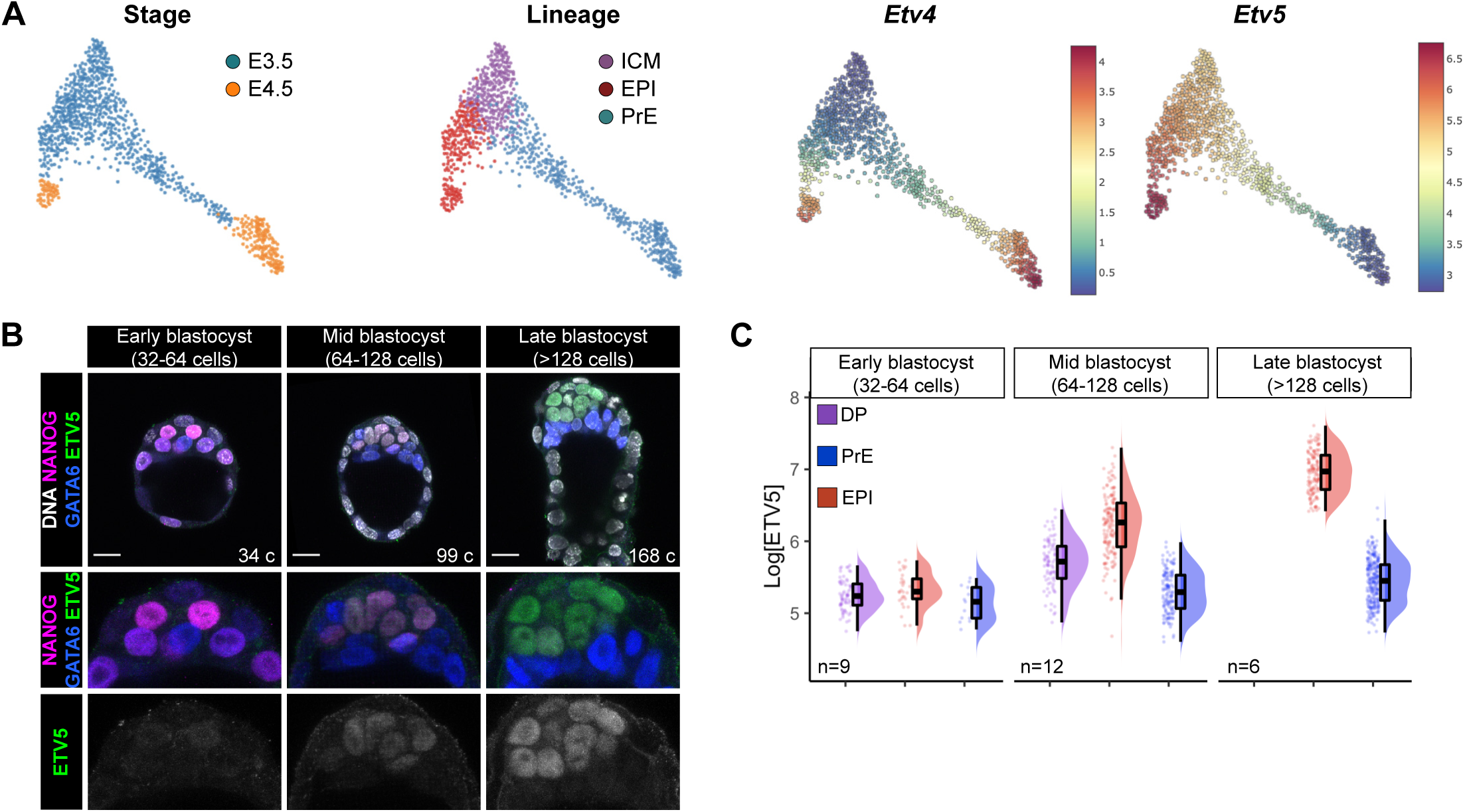
*Etv4* and *Etv5* expression during mouse embryonic development. **(A)** Single-cell RNAseq from (Nowotschin et al 2019) showing *Etv4* and *Etv5* mRNA expression at E3.5 and E4.5 in inner cell mass (ICM), epiblast (EPI) and primitive endoderm (PrE) lineages. **(B)** Confocal images of immunofluorescence immunostaining of NANOG, GATA6 and ETV5 in blastocyst stage mouse embryos. Cell numbers (c) indicated. Scale bars 20 μm **(C)** Quantification of ETV5 levels from (B). Double positive (DP; NANOG+GATA6), epiblast (EPI; NANOG+) and primitive endoderm (PrE; GATA6+)

Next, we assessed protein expression at pre-implantation stages. We were unable to detect the ETV4 protein at pre-implantation stages, likely due to the lack of a good commercially available antibody, however ETV5 protein was readily detectable. We performed immunofluorescence staining for ETV5, NANOG, GATA6, and quantified nuclear fluorescence intensity to categorise cell lineages (Supplemental Fig. 1A,B), as described previously (Saiz et al., 2016). In early blastocyst (32-64 cells) stage embryos, ETV5 protein was not expressed. At mid-blastocyst (64-128 cells) stage, low levels of ETV5 protein were detected in uncommitted ICM progenitor cells (NANOG+GATA6+), and at higher levels in epiblast cells (NANOG+GATA6-) (Fig. 1B,C). The highest levels of ETV5 protein were detected in the epiblast cells (NANOG+GATA6- or NANOG-GATA6-) of late blastocyst stage embryos (Fig. 1B,C). ETV5 protein was not present in the primitive endoderm (NANOG- GATA6+) or outer trophectoderm cells.

Further to this, we assessed early post-implantation embryos for ETV5 protein expression. We performed immunofluorescence analysis for ETV5, GATA6 (visceral endoderm marker), and SOX2 (epiblast and extra-embryonic ectoderm marker). At the egg cylinder stage (E5.5), ETV5 protein is lowly expressed within the nuclei of the epiblast and in the distal-most part of the extra-embryonic ectoderm adjacent to the epiblast, and absent in the visceral endoderm (Supplementary Fig. 1C). At mid-gastrulation stage (E6.5), ETV5 was expressed heterogeneously in the epiblast and in the nascent mesoderm (Supplementary Fig. 1C).

Taken together, the transcript and protein profiling reveal that *Etv5* is first expressed in ICM progenitors, and then specifically upregulated in epiblast cells, where expression is maintained in these pluripotent cells through peri- and post-implantation stages.

### Loss of *Etv5* compromises the formation of primitive endoderm

To determine if ETVs might act as downstream effectors of FGF signalling in pre-implantation development, we next wanted to assess the role ETVs in blastocyst formation. Therefore, we analysed cell lineages upon loss of one, or both, *Etv4* (Livet et al., 2002) and *Etv5 (Zhang et al., 2009)* genes. We collected embryos from *Etv4^+/-^;Etv5^+/-^* inter-cross matings at early to late blastocyst stage, and assessed the number of cells in each lineage by quantitative immunofluorescence staining for CDX2 (trophectoderm), NANOG (epiblast) and GATA6 (PrE, Supplemental Fig. 2A) as described earlier. Uncommitted ICM progenitor cells are specified asynchronously from early blastocyst stage to epiblast and primitive endoerm, so that by the late-blastocyst (>128 cells) stage there is a stereotyped proportion of 40% epiblast cells and 60% primitive endoderm within the inner cell mass (Saiz et al., 2016).

From the *Etv4^+/-^;Etv5^+/-^* inter-cross progeny, wild-type embryos, and embryos with at least one intact copy of *Etv5*, specified the correct ratio of epiblast cells (Fig. 1A,B,C). Therefore, loss of *Etv4* alone does not alter the epiblast:PrE ratio. Complete loss of *Etv5* (embryos with the genotypes *Etv4^+/+^;Etv5^-/-^, Etv4^+/-^;Etv5^-/-^, and Etv4^-/-^;Etv5^-/-^*), by contrast, had a marked effect on the ratio of epiblast:PrE at both mid-blastocyst and late blastocyst stage (Fig. 1A,B,C). *Etv5^-/-^* embryos, regardless of *Etv4* genotype, had a reduction in the proportion of primitive endoderm cells (57-63% epiblast: 37-43% PrE) by the late blastocyst stage (Fig. 2C). *Etv5^-/^*^-^ embryos exhibited a range in the fraction of primitive endoderm cells in the ICM, with some embryos having fewer percentage of primitive endoderm cells, while others exhibit a complete loss (Fig. 2A,B, Supplemental Fig. 2B). The *Etv5^-/-^* embryos phenocopy *Fgfr1^-/-^* embryos (Kang et al., 2017; Molotkov et al., 2017), suggesting a heirachecical relationship of *Fgfr1* and *Etv5* in relaying FGF signalling to specify primitive endoderm.

**Figure 2:**
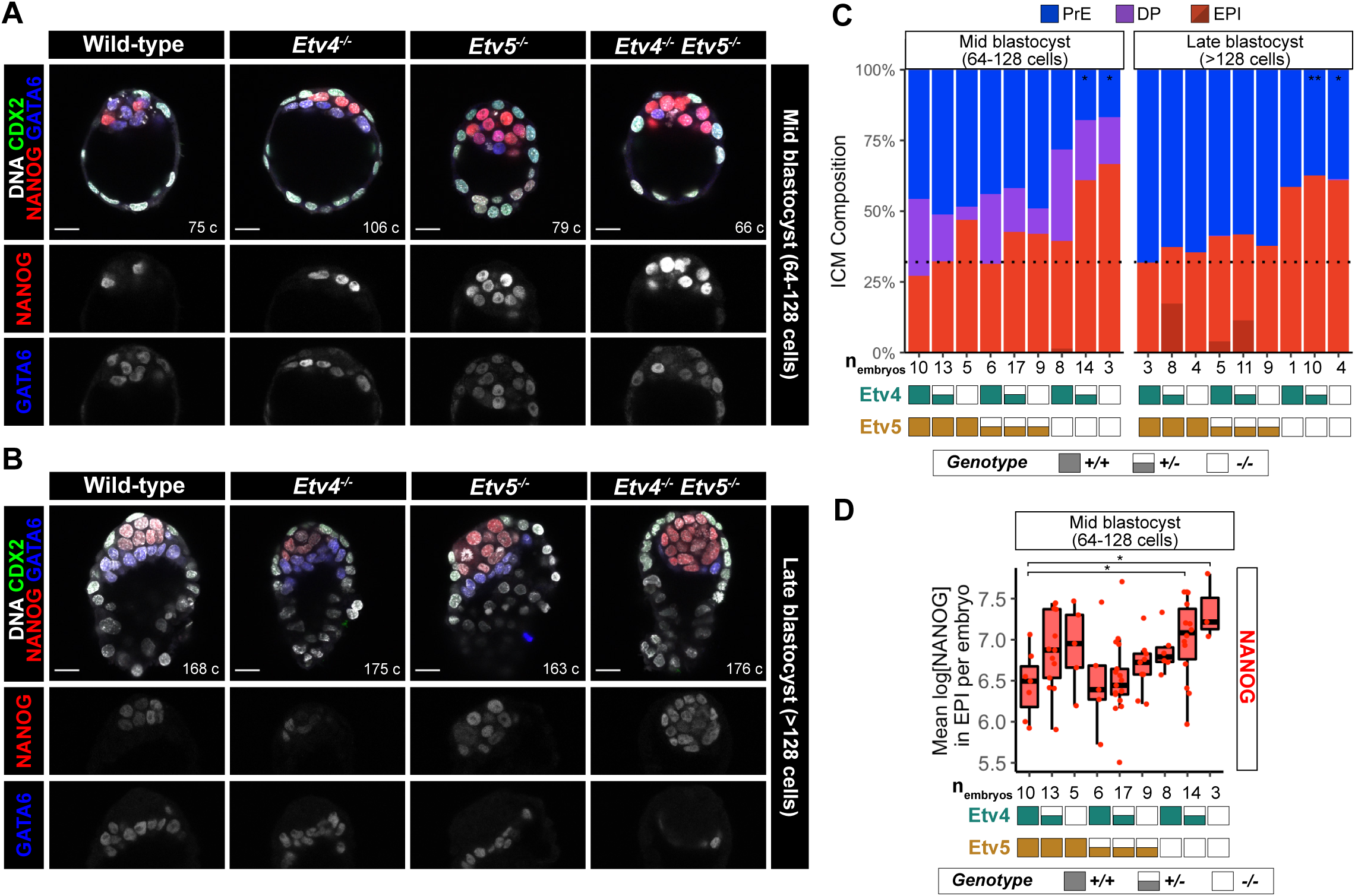
Loss of *Etv5* compromises the formation of primitive endoderm. **(A-B)**Confocal images of immunofluorescence staining of CDX2, NANOG and GATA6 in an *Etv4;Etv5* allelic series of embryos at mid-**(A)** and late- **(B)** blastocyst stages. Scale bars 20 μm **(C)** Quantification of inner cell mass (ICM) lineage composition in an allelic series of *Etv4;Etv5* mutant embryos shown in (A-B). Double positive (DP; NANOG+GATA6+), primitive endoderm (PrE; NANOG-;GATA6+), epiblast (EPI; NANOG+;GATA6-light red, NANOG-;GATA6-dark red). Black dotted line represents wild-type ratio of EPI:PrE at late blastocyst stage. T-test of PrE numbers compared to wild-type, * p < 0.05, ** p < 0.01, otherwise comparison not significant. *Etv4^+/+^;Etv5^-/-^* at late-blastocyst stage did not have sufficient n number to performstatistical test. **(D)** Quantification of NANOG levels in epiblast cells in wild-type embryos and an allelic series of *Etv4;Etv5* mutant embryos shown in (A) at mid-blastocyst stage. T-test of mean NANOG levels per embryo compared to wild-type * p < 0.05, otherwise comparison not significant.

The reduction in the proportion of primitive endoderm cells in *Etv5* null embryos was driven by both an increase in epiblast cell numbers and a decrease in primitive endoderm cell numbers, without changes in total cell counts (Supplemental Fig. 2C). The compensatory increase in epiblast versus primitive endoerm cell numbers in *Etv5* null embryos suggested that uncommitted cells were preferentially specified to epiblast. While the similarity in the number of DP cells in single *Etv5*^-/-^ and *Etv4^-/-^*, and double *Etv4^-/-^;Etv5^-/-^* knockout embryos, and wild-type embryos indicated timely specification of uncommitted progenitors for all embryo genotypes (Fig. 2C). These data, combined with the comparable number of ICM cells between all genotypes (Supplemental Fig. 2D), argue against a delay in specification, or the selective loss of primitive endoerm cells.

Although epiblast specification did not appear to be negatively affected in any of the mutant embryos, we wanted to determine if the epiblast lineage was being properly maintained after specification. Declining NANOG levels are indicative of the exit of naïve pluripotency as the epiblast undergoes maturation prior to implantation, and is dependent on FGF signalling (Kang et al., 2017; Molotkov et al., 2017; Nichols et al., 2009). NANOG levels were elevated in *Etv4^-/-^* and *Etv5^-/-^* single knockout embryos, when compared with wild-type embryos at the late-blastocyst stage (Fig. 2D). This unphysiologically elevated level of NANOG was further compounded in double knockout embryos (Fig. 2D), with NANOG levels significantly higher in *Etv4^+/-^;Etv5^-/-^*and *Etv4^-/-^;Etv5^-/-^* compared to wild-type embryos, suggesting that both ETV factors play complementary and synergistic roles in the maturation of the epiblast lineage.

All together, these data suggest *Etv5*, but not *Etv4*, is required for balancing the specification of uncommitted ICM cells towards primitive endoerm and away from epiblast cells, to achieve the robust and stereotyped tissue proportions of the blastocyst.

### Mechanism of *Etv5* action on inner cell mass cell fate decision

Given the phenotypic similarity of *Etv5* mutants to mutants with reduced FGF signalling activity (e.g. *Fgf4^+/-^* and *Fgfr1^-/-^* embryos), we reasoned that FGF signalling may be disrupted or dampened in *Etv5* mutant embryos. *Etv5* is expressed at intermediate levels in uncommitted ICM cells, and upregulated in epiblast cells, mirroring the expression of *Fgf4* (Fig. 1, Supplemental Fig. 1B, (Nowotschin et al., 2019). Hypothesising that *Etv5* may regulate *Fgf4* transcription, we analysed ChIP-seq data of ETV5 binding (Kalkan et al., 2019) in mouse embryonic stem cells (mESC) cultured in 2i (naïve pluripotent conditions), and 16h after transfer to N2B27 (to induce transition out of the naïve pluripotent state). ETV5, a transcriptional activator, is enriched at the upstream promoter region of *Fgf4* in naïve and transitioning mESCs (Fig. 3A), suggesting direct regulation in epiblast cells *in vivo* as well.

**Figure 3:**
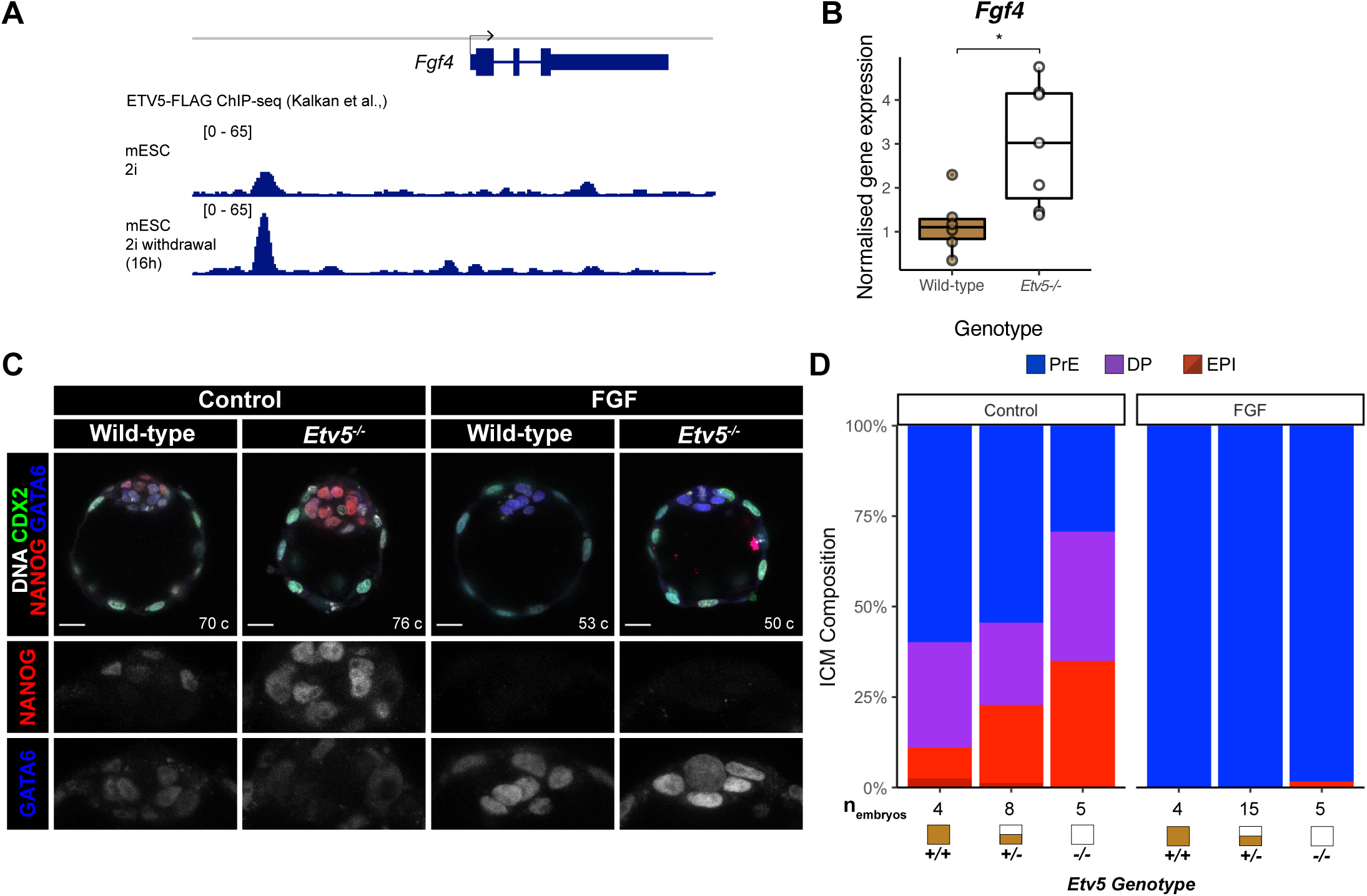
Mechanism of *Etv5* action on inner cell mass cell fate decision. **(A)**ETV5 binding to the *Fgf4* locus in mouse embryonic stem cells (mESC) in naïve pluriptency conditions (2i) and upon naïve pluropotency exit (16 hours after 2i withdrawal). ChIP-seq data (Kalkan et al 2019). **(B)** Expression of *Fgf4* in wild-type and *Etv5^-/-^* late-stage blastocyst by qPCR. T-test, * p < 0.05. **(C)** Confocal images of immunofluorescence staining of CDX2, NANOG and GATA6 in wild-type and *Etv5^-/-^* mutant embryos treated with or without 1μg/ml FGF4 + 1μg/ml Heparin from E2.5 + 48 hours. Scale bars 20 μm. **(D)** Quantification of inner cell mass (ICM) lineage composition in treated embryos (C). Double positive (DP; NANOG+GATA6+), primitive endoderm (PrE; NANOG-;GATA6+), epiblast (EPI; NANOG+;GATA6-light red, NANOG-;GATA6-dark red).

To test this hypothesis, we analysed *Fgf4* expression after loss of *Etv5* in embryos. Quantitative real-time PCR (qRT-PCR) of E4.5 whole blastocysts demonstrated that *Fgf4* transcripts are elevated in *Etv5^-/-^*embryos compared with wild-type (Fig. 3B, fold change = 1.59, P = 0.01). Therefore, in embryos, *Etv5* is dispensable for *Fgf4* expression, which is sustained in *Etv5^-/-^* embryos with higher *Fgf4* expression correlating with increased number of epiblast cells. Thus, loss of primitive endoderm in the *Etv5* mutants is not caused by non-cell autonomous effect of limited FGF4 availability.

Given the dampened response to FGF signalling in the *Etv5^-/-^*embryos, we attempted to rescue the reduction in primitive endoderm numbers with excess exogenous FGF ligand. We treated wild-type, *Etv5^+/-^*, and *Etv5^-/-^* embryos with a saturating dose of FGF4 (1μg/ml) and Heparin (1μg/ml) from E2.5 (8-16cell stage) for 48 hours. After FGF treatment, *Etv5^-/-^*embryos consisted of GATA6+NANOG-primitive endoderm cells throughout their ICMs, similar to wild-type and heterozygote embryos (Fig. 3C,D). These experiments demonstrate that high, non-physiological levels of exogenous FGF4 can rescue the specification of the primitive endoerm in *Etv5^-/-^* mutants. In conclusion, these experiments show that while primitive endoderm specification can be rescued with exogenous FGF4, under physiological conditions, *Etv5* plays a crucial cell-autonomous role in tuning sensitivity of ICM cells to the FGF4 signal during primitive endoderm specification.

### Loss of *Etv4/5* causes a delay in the progression of pluripotency

We hypothesised that ETV factors may play a role in epiblast maturation, as suggested by the elevated NANOG levels observed in mutant embryos (Fig. 2D). To probe the role of *Etv4* and *Etv5* in pluripotency exit further, we assessed additional markers of pluripotency at the late blastocyst stage. Embryos from *Etv4^+/-^;Etv5^+/-^*inter-crosses were stained for a core pluripotency marker (SOX2), naïve marker (KLF4), and formative/primed marker (OTX2) (Fig. 4A). While SOX2 and KLF4 in the epiblast were expressed at similar levels across all genotypes (Supplemental Fig. 4A, B), epiblast OTX2 expression was significantly reduced in embryos lacking *Etv5* (Fig. 4B, C). Analysis of published Etv5 chromatin binding in mESC (Kalkan et al., 2019) shows strong binding at a downstream enhancer region (Fig. 4D).

**Figure 4:**
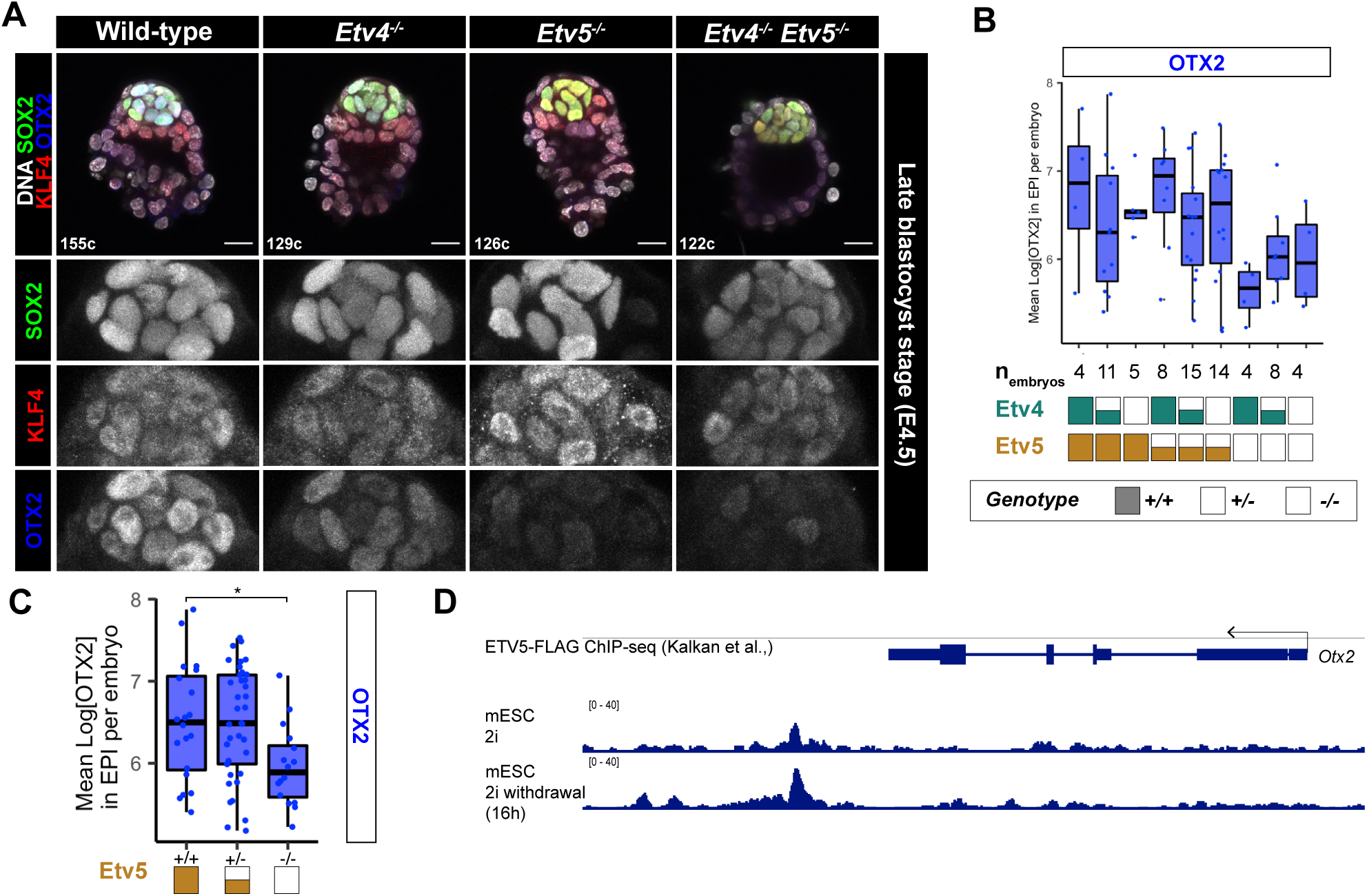
Loss of *Etv4/5* causes a delay in the progression of pluripotency. **(A)**Confocal images of immunofluorescence staining of SOX2, OTX2 and KLF4 in an allelic series of *Etv4;Etv5* mutant embryos at the late blastocyst stage. **(B-C)** Quanitifcation of primed marker OTX2 from (A) by individual genotypes **(B)** or grouped by *Etv5* genotype **(C)**. T-test of mean OTX2 levels in epiblast cells per embryo, compared with wild-type, * p < 0.05, otherwise comparison was not significant. **(D)** ChIP-seq of ETV5 binding to the *Otx2* locus in mouse naïve ESC (2i) and after pluripotency exit (2i withdrawal). Data from (Kalkan et al., 2019).

Together, these data indicate that during pre-implantation development *Etv5* is required for timely exit of pluripotency, likely through direct regulation of *Otx2*, and indirect regulation of *Nanog* by an as yet unknown mechanism.

We then assessed pluripotency at early post-implantation stages at E5.5. In wild-type, *Etv4*, and *Etv5* null mutants embryos, the epiblast cavitated and retained expression of the core pluripotency factor SOX2 (Supplemental Fig. 4C). Interestingly, OTX2 was expressed in both the visceral endoderm and the epiblast at this stage, indicating that ETV factors are not necessary for upregulation of primed pluripotency factors at early post-implantation stage.

We then tested the requirements of *Etv4* and *Etv5* for pluripotency in ESCs. We differentiated double knockout *Etv4^-/-^;Etv5^-/-^*ESCs (Lu et al., 2009) to epiblast-like cells (EpiLC) representative of the early post-implantation epiblast, which required exogenous FGF2 and Activin. After two days of EpiLC differentiation, both wild-type and double knockout cells maintained core pluripotency transcription factors (OCT4, SOX2), downregulated naïve markers (NANOG, KLF4) and upregulated primed markers (Supplemental Fig. 4D,E). Indicating, that loss of *Etv4/5* is not sufficient to completely block naïve pluripotency exit, in agreement with *in vivo* early post-implantation embryo data (Supplemental Fig. 4C). However, given the failure in downregulation of NANOG and upregulation of OTX2 at pre-implantation stages, ETVs likely control the timely exit of naïve pluripotency in the embryo, this delay has been similarly shown in ESCs (Kalkan et al., 2019).

### Compound *Etv4/5* mutants display developmental delay and anterior visceral endoderm migration defects

We next looked at later embryonic time-points to determine if the delayed naïve pluripotency exit impacted later stages of development. At E6.5, prior to gastrulation, wild-type and *Etv4* mutant embryos expressed high levels of OTX2 in the anterior visceral endoderm (AVE), and NANOG expression was confined to the proximal posterior region of the epiblast, marking the primitive streak (Fig. 5A). In *Etv5* null, and compound mutants, the distal visceral endoderm (DVE) marked by high OTX2 failed to migrate anteriorly (Fig. 5A, left panel arrow heads). In single *Etv5* mutants, this resulted in a failure to properly position the anterior-posterior axis, as evidenced by the ∼45 degree rotation of the NANOG expression domain, marking the primitive streak. Double knockout embryos at E6.5 were overall smaller compared to E5.5 wild-type embryos. The extra-embryonic ectoderm region was noticeably reduced in size, with a disordered morphology of the visceral endoderm epithelium (Fig. 5A, right panel arrowhead). NANOG expression was elevated throughout the epiblast, indicating that either the entire epiblast had failed to exit naïve pluripotency, or the presence of an expanded primitive streak region.

**Figure 5:**
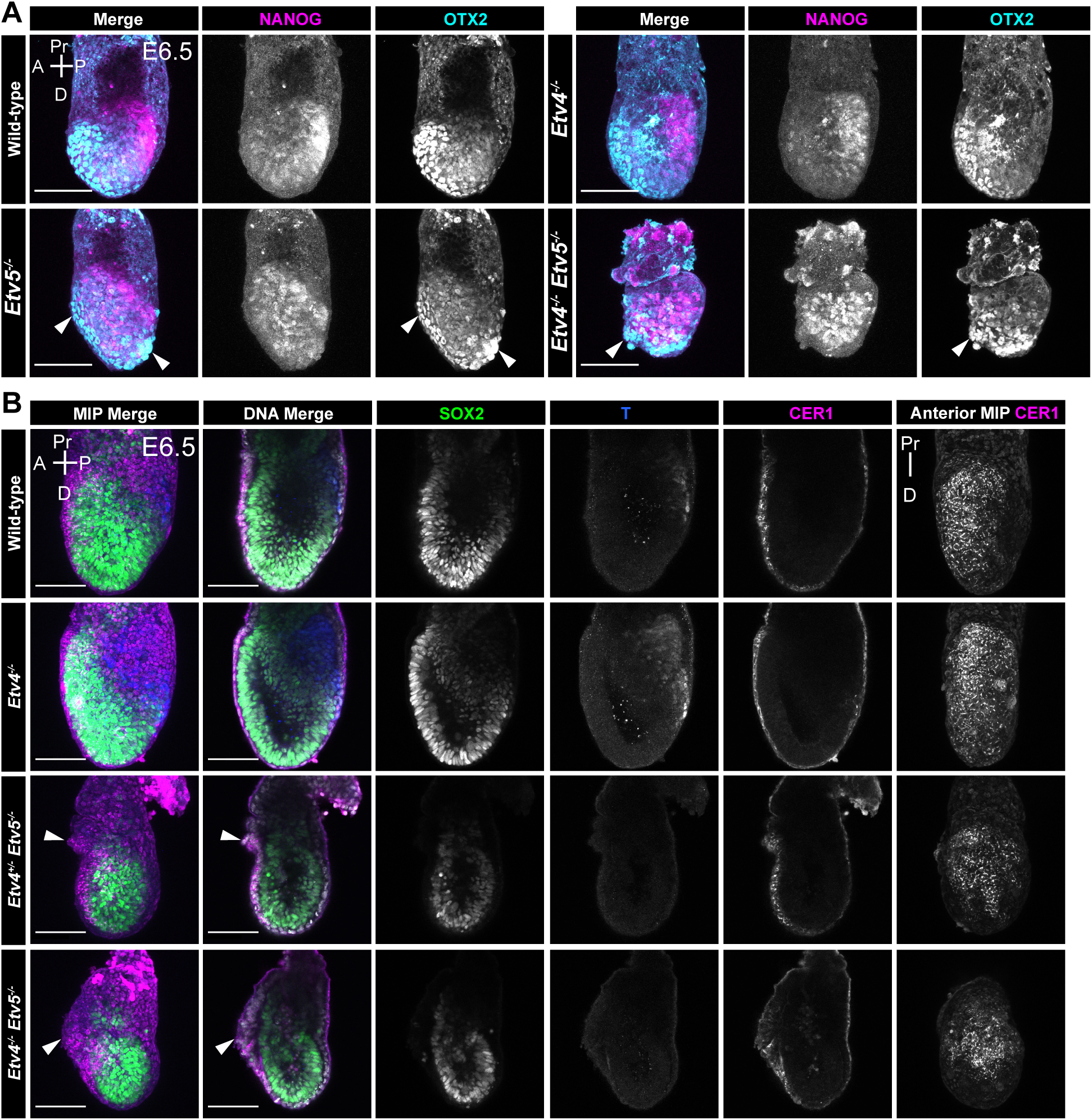
Compound *Etv4/5* mutants have developmental delay and anterior visceral endoderm migration defects. **(A)**Confocal max intensity projection images of an allelic series of *Etv4;Etv5* embryos at mid-streak gastrulation stages, embryonic day (E) 6.5, immunostained for NANOG and OTX2. Arrowheads show abnormal anterior/distal visceral endoderm migration and/or morphology. **(B)** Confocal images of an allelic series of *Etv4;Etv5* embryos at mid-streak gastrulation stages, embryonic day (E) 6.5 immunostained for SOX2, T, and CER1. Arrowheads show abnormal anterior visceral endoderm migration and morphology. MIP = max intensity projection. A = Anterior, P= Posterior. Pr = Proximal. D = Distal. Scale bars 100μm.

To determine if mutant embryos were specifying the AVE correctly and able to initiate gastrulation, we stained for an AVE marker, CER1, and mesoderm marker, T (Brachyury). In wild-type and *Etv4* null embryos, CER1 was localised to the anterior region of the embryo, extending from the embryonic-extraembryonic junction to the distal tip (Fig. 5B), while T was localised to the posterior epiblast of marking the nascent mesoderm. However, in single *Etv5* and double *Etv4;Etv5* homozygous mutant embryos, while CER1 was expressed, indicating that the AVE had been specified, the migration of the AVE appeared to be disrupted, delayed (Fig. 5B, arrowheads), or arrested at the distal tip (Supplemental Fig. 5A). The AVE, as evidenced by CER1 expression could sometimes be seen extending beyond the embryonic-extraembryonic junction, indicting an over migration of the AVE. The CER1 domain size was noticeably reduced relative to the size of the embryos. Again, abnormal thickening, and disordered DVE/AVE epithelial morphology was observed in these mutants (Fig. 5B arrowhead). Finally, mutants were devoid of T+ cells, indicating a failure in gastrulation and/or specification of mesoderm cell types.

The AVE migration phenotype and reduced embryo size was more severe and pronounced in compound mutant embryos *Etv4^+/-^;Etv5^-/-^* and *Etv4^-/-^;Etv5^-/-^*when compared with single homozygous *Etv4^+/+^;Etv5^-/-^*embryos. So, while *Etv4* single mutants did not have an overt phenotype, partial or complete loss of *Etv4* did increase the severity of the *Etv5* phenotype, suggesting that these factors may partially compensate, or have overlapping roles in the early post-implantation epiblast.

By late-gastrulation (E7.5) *Etv5* and double homozygous null embryos were severely developmentally retarded with a failure in gastrulation, as indicated by the absence of the primitive streak marker, T (Supplemental Fig. 5B). Epiblast morphology was grossly abnormal, with ruffles and folds to the epithelium. As a consequence, *Etv5* homozygous mutant pups were not recovered from *Etv4;Etv5* heterozygous intercrosses, and were drastically under-represented in *Etv5* heterozygous intercrosses (Supplemental Fig. 5C,D). *Etv4^-/-^;Etv5^+/-^* pups were also not recovered, further suggesting a combinatorial role for ETV factors during embryonic development. All together, these data demonstrate that embryos of *Etv4;Etv5* allelic series have increasingly severe phenotypes with loss of ETV gene dosage at early postimplantation stages exhibiting developmental delay and AVE migration defects which compromise embryonic development.

## Discussion

Here, we have shown that *Etv5^-/-^* blastocysts have a reduction in primitive endoderm specification. This phenotype bears a striking similarity to that of *Fgfr1*^-/-^ and *Fgf4*^+/-^ embryos (Brewer et al., 2015; Kang et al., 2017; Kang et al., 2013). Given that *Fgf4* transcripts are not diminished in *Etv5* mutants, the phenocopying of low dosage FGF mutants cannot be due to a decreased ligand availably. Our findings support the hypothesis that *Etv5* relays FGF signalling activity in uncommitted ICM cells, as in other developmental contexts (Herriges et al., 2015; Zhang et al., 2009). The direct involvement of *Etv5* in initiating the primitive endoderm program is further supported by its ability to upregulate multiple endoderm genes (including *Sox7* and *Sox17*) when overexpressed in mouse ESCs (Correa-Cerro et al., 2011). However, FGF hyperstimulation can rescue primitive endoderm cell numbers in *Etv5*-deficient embryos, suggesting that under these non-physiological conditions, related ETS family members such as *Etv1* and *Etv4* may compensate for loss of *Etv5* function.

NANOG, a hallmark marker of mouse naïve pluripotency, is downregulated in the epiblast prior to implantation in an FGF/ERK dependent manner (Nichols et al., 2009). In both *Etv4^-/-^* and *Etv5^-/-^*blastocysts, elevated NANOG levels reveal a failure to exit naïve pluripotency. *Etv4* and *Etv5*, have been implicated in regulating pluripotency *in vitro*. For example, *Etv5* was shown to promote MET during iPSC reprogramming (Zhang et al., 2018). In addition, these ETV factors appear to regulate ESC proliferation, but there is conflicting evidence as to whether they promote or repress epiblast-like fate during differentiation (Akagi et al., 2015; Zhang et al., 2018). Triple knock-out of *Etv5, Rbpj* and *Tcf3* in ESC can maintain naïve pluripotent state in absence of 2i (Kalkan et al., 2019), implicating these factors in dissolution of the naïve pluripotency transcription factor network. Given that ETV5 does not directly bind to the *Nanog* locus in ESC (Kalkan et al., 2019), it seems likely that this regulation by ETV factors *in vivo* is indirect. Instead, ETVs may activate the primed pluripotency network, which then, in turn, repress the naïve state. Consistent with such a model, the primed pluripotency marker, OTX2, fails to be upregulated in *Etv5* mutant blastocysts. ETV5 has been shown to directly bind an *Otx2* enhancer in mESCs (Kalkan et al., 2019), suggesting it is a direct target *in vivo*. As *Otx2* and *Nanog* are antagonistic (Acampora et al., 2017; Acampora et al., 2016), the role of ETV in epiblast cells may be to turn on *Otx2*, and consequently reduce *Nanog* levels thereby promoting the exit from naïve pluripotency.

OTX2, a marker of primed pluripotency, is normally expressed in the epiblast from late blastocyst stage (E4.5) through early post-implantation. In *Etv5*^-/-^ embryos however, OTX2 is absent at E4.5 but eventually expressed by E5.5, indicating a delayed exit from naïve pluripotency. This delay is consistent with our observation of elevated NANOG levels in *Etv5* mutant blastocysts. Fruthermore, this is further supported by time-course analysis of *Etv5^-/-^* ESC differentiation showing delayed naïve pluripotency exit (Kalkan et al., 2019).Together, these findings highlight the role for the ETVs in regulating the robustness and timely exit of naïve pluripotency.

Timely, Nodal-dependent maturation of the epiblast at early post-implantation stages is required for the co-ordinated and spatial patterning of the embryo and initiation of gastrulation (Huang et al., 2017; Zang et al., 2022). Given that *Etv4* and *Etv5* are not expressed in the visceral endoderm, we hypothesize that the delayed epiblast maturation observed in *Etv5* and *Etv4;Etv5* compound mutants leads to the later AVE development defects. In the *Etv5* and *Etv4;Etv5* compound mutants there is a reduction in the number and migration CER1 expressing cells, with disordered discontinuous AVE and DVE populations, and ectopic protrusions. These phenotypes are reminiscent of mutants affecting AVE development, including mutants in Nodal signalling pathway components (Stower and Srinivas, 2014). The most severely affected *Etv4^-/-^;Etv5^-/-^* embryos either fail to migrate the AVE/DVE, or the AVE over-migrates beyond the embryonic-extraembryonic boundary, and exhibit a disordered multi-layered epithelium, reminiscent of the two classes of *Lefty1* mutants (a Nodal antagonist) (Trichas et al., 2011). Our research suggests that FGF, in conjunction with Nodal, plays a role in epiblast development and its interaction with the visceral endoderm, which is crucial for establishing the anterior-posterior axis.

All together, our results demonstrate the successive roles of ETS factors *Etv4* and *Etv5* as FGF signaling effectors in early lineage specification and embryonic body plan establishment, increasing our understanding of the molecular mechanisms of mammalian development.

## Materials and Methods

### Immunofluorescence

Pre-implantation embryo immunofluorescence was carried out as previously described (Saiz et al., 2016). Briefly, embryos were fixed for 10 min at room temperature in 4% PFA. Fixed blastocysts were washed in PBX; 0.1% Triton X-100 (Sigma-Aldrich) in PBS, permeabilised for 5 min in a solution of 0.5% Triton X-100, 100 mM glycine in PBS and then washed in PBX for 5 min. Embryos were blocked in blocking buffer: 2% horse serum in PBS, for 40 min at room temperature, followed by incubation overnight at 4°C with primary antibodies diluted in blocking buffer (see antibody list below). The next day, embryos were washed three times in PBX, incubated in blocking buffer for 40 min at room temperature before a 1 h incubation with secondary antibodies at 4°C. Embryos were then washed in PBX and incubated in 5 μg/ml Hoechst in PBS for at least 30 min prior to imaging.

For post-implantation stages, embryos were fixed for 30 min at room temperature in 4% PFA. Then, fixed embryos were washed in PBX, permeabilised in 0.5% Triton-X in PBS for 30 min and then washed three times in PBX. Embryos were then incubated in blocking buffer, 5% donkey serum and 0.2% BSA in PBX for 2hrs at RT, followed by incubation overnight at 4°C with primary antibodies diluted in blocking buffer. The next day, embryos were washed in PBX, followed by a second blocking step for at least 2hrs at room temperate, and incubation with secondary antibodies in blocking buffer overnight at 4 °C. After antibody staiing, embryos were washed in PBX, and incubated with 5 μg/ml Hoechst in PBX for a minimum of 2 hours to visualise DNA prior to imaging.

### Cell lines

ESC lines used in the study were wild-type and *Etv4^-/-^;Etv5^-/-^*cells (Lu et al., 2009). ESCs were maintained on 0.1% gelatin (Millipore, 104070) coated tissue-culture grade plates in a humidified 37°C incubator with 5% CO2. ESCs were grown in DMEM (Life Technologies, 11995073), supplemented with 2mM L-glutamine (Life technologies, 25030164), 0.1mM MEM NEAA (Life technologies, 11140-050), 1mM sodium pyruvate (Life technologies, 11360070), 100U/ml penicillin and 100mg/ml streptomycin (Life technologies, 15140163) , 0.1mM 2-mercaptoethanol (Life technologies, 21985023), 15% FBS (VWR, 97068-085), and 1000U/ml LIF (prepared in house). ESC were differentiated to EpiLC as previously described (Hayashi et al., 2011). Briefly, ESCs were seeded 2.5 x 10^4^ cells/cm^2^ in EpiLC medium onto fibronectin (16μg/ml, Millipore, FC010) coated 8-well IBIDI plates. EpiLC medium was comprised of N2B27 Medium containing 20 ng/ml ACTIVIN A (Peprotech 120-14P), 12 ng/ml FGF2 (R&D, 233-FB-025), and 1% KSR (Thermo 10828028)

### Antibodies

Antibodies:
|  |  |
| --- | --- |
| ETV5 | Rabbit Proteintech 13011-1-AP 1:100 |
| GATA6 | Goat R&D AF1700 1:100 |
| NANOG | Rabbit Reprocell REC-RCAB0002PF 1:500<br>Rat e-biosciences 14-5761 1:100 |
| SOX2 | Rat e-biosciences 14-9811-82 1:200 |
| KLF4 | Rabbit Cell Signalling 4038P 1:100 |
| OTX2 | Goat R&D AF1979 1:500 |
| CER1 | Goat R&D AF1986 1:300 |
| T | Goat R&D systems AF2085 1:100<br>Rabbit Abcam ab209665 1:100 |

### Mice & Genotyping

*Etv4^tm1Arbr^* (Livet et al., 2002) and *Etv5^tm1.1Xsun^*(Zhang et al., 2009) knockout alleles (abbreviated to *Etv4^-^* or *Etv5^-^*) were maintained as double and single heterozygous mouse lines on a CD1 background. Mice, whole blastocysts, whole embryos or trophectoderm cells were genotyped by PCR. Three primer PCR was used to genoptype *Etv5* samples, as follows: F 5’-CTCGCAGAGGACAAGGTAGTGAC-3’ R_WT 5’-GTGTGCACGACATGTTCAAGG-3’ R_KO 5’-CCAGCATCGTACAAAACAAGAG-3’, generating a wild-type band 270 bp and a knock-out band at 374 bp. For genotyping *Etv4* samples, the primers Etv4_WT_F 5’-TCTGGACCCTCTCCAGGTGATG-3’ and Etv4_WT_R 5’-CCACCAGAAACTGCCACAGTTG-3’ generating a wild-type band of 501bp, and LacZ_F 5’-CATCCACGCGCGCGTACATC-3’ LacZ_R 5’-CCGAACCATCCGCTGTGGTAC-3’ generating a knock-out band, amplifying the LacZ cassette of 360 bp.

### FGF embryo treatment

Embryos for this study were obtained from natural matings between *Etv5* male and female heterozygotes. The sex of embryos was not determined. E2.5 morulae were flushed from oviducts with flushing holding medium (FHM, Millipore) as described (Behringer et al., 2014). Embryos within litters were randomly assigned in even sized groups for control and exogenous FGF treatment. Control: KSOM (MR-121-D, Sigma) and FGF stimulation: 1μg/ml FGF4 (R&D Systems) and 1μg/ml Heparin (Sigma) in KSOM. Medium was equilibrated 30 mins prior to culture to reach the correct temperature and pH. Embryos were cultured in groups in droplets of medium in 35mm dishes (approximately 1μl/embryo) overlaid with mineral oil (Sigma), for 48 hours in total in a humidified incubator at 37°C with 5% CO2. After 24 hours (E3.5) the zona pellucidae were removed by brief incubation in acid Tyrode’s solution (Sigma), washed three times in their respective culture medium, and then cultured in fresh droplets of medium. Embryos were assay for lineage markers by immunofluorescence at the end of 48hr culture.

### Single-embryo qPCR

E4.5 stage embryos from *Etv5* intercrosses were prepared for genotyping and qPCR as previously described (Kang et al., 2017; Morgani et al., 2018), using CellsDirect One-Step kit in accordance with the manufacturer’s instructions. Blastocyst were first washed in PBS, then incubated in 0.5% trypsin for 3 mins at 37°C. Using a glass capillary, a small number of mural TE cells were removed for PCR genotyping, then each blastocyst was added to 5 µl of 2 × Reaction Mix (Invitrogen, CellsDirect One-Step qRT-PCR Kit) and snap-frozen on dry ice and stored at -80°C until processing.

For cDNA and target specific pre-amplification 5μl of a reverse transcription/pre-amp mix was added to each blastocyst lysate. For each sample this comprised of 0.2μl SuperScript II RT/Platinum Taq mix (Invitrogen), 2.5μl TaqMan assay (pooled assay mix with a concentration of 0.2x for each probe, detailed in supp. Table X), 2.3μl RNase-free H2O. To perform combined cell lysis, cDNA synthesis and pre-amplification of specific targets, samples were incubated at 50 °C for 20 min, 95 °C for 2 min, followed by 18 cycles of 95 °C for 15 sec then 60 °C for 4 min, in a T100 Thermal Cycler (Bio-Rad).

Blastocyst cDNA was diluted 1 in 5 by adding 40μl H20 to the 10μl cDNA to a total of 50μl. To assay the amount of mRNA, each qPCR reaction was set up in duplicate in 96-well plates (Applied Biosystems 4306737) overlaid with MicroAmp clear adhesive film (Applied Biosystems 4306311). For each qPCR reaction the mix was as follows: 7.5μl TaqMan Universal PCR Master Mix, 0.75μl TaqMan Assay, 5.25μl RNase free H2O. 1.5μl diluted sample cDNA was then added to each reaction, to a total of 15μl. Real-time PCR was then carried out in a QuantStudio 7 Flex System (Applied Biosystems). Gene expression was calculated as 2^ΔΔCt^. Target gene (FGF) expression was calibrated to the arithmetic mean of the wild-type samples, and normalised to two reference genes’ expression within each sample, *Gapdh* and *Actb,* using the geometric mean (Vandesompele et al., 2002).

## Acknowledgements

We thank the members of the Hadjantonakis and Niakan labs for helpful discussions and comments on the manuscript. We thank the Francis Crick Institute’s Genomics Equipment Park.

## Supplementary figure legends

**Supplemental figure 1: *Etv4* and *Etv5* expression during mouse embryonic development (related to** Figure 1)

**(A)** Quantification of NANOG and GATA6 levels in blastocyst (from Figure 1B). Clustering into double positive (DP; NANOG+;GATA6+), epiblast (EPI; NANOG+;GATA6-) and primitive endoderm (PrE; NANOG-;GATA6+)

**(B)** Heatmap of ETV5 levels in blastocyst compared to NANOG and GATA6 levels (from Figure 1B)

**(C)** Confocal images of immunofluorescence immunostaining of SOX2, GATA6 and ETV5 in postimplantation stage stage mouse embryos. Scale bars 50μm. Anterior (A); posterior (P); proximal (Pr); distal (D); extraembryonic-ectoderm (ExE); epiblast (Epi); PS (Primitive streak).

**Supplemental figure 2: Loss of *Etv5* compromises the formation of PrE (related to figure 2)**

**(A)** Quantification of NANOG and GATA6 levels in an *Etv4;Etv5* allelic series of mutant embryos (from Figure 2A,B). Clustering into double positive (DP; NANOG+;GATA6+), primitive endoderm (PrE; NANOG-;GATA6+), epiblast (EPI; NANOG+;GATA6-light red, NANOG-;GATA6-dark red).

**(B)** Quantification of inner cell mass (ICM) lineage composition in an allelic series of *Etv4;Etv5* mutant embryos. Individual embryos shown and ordered by ascending cell number. Dotted line repressends mean wild-type EPI:PrE composition by late blastocyst stage.

**(C)** Total number of cells per embryo in EPI and PrE lineages

**(D)** Total number of cells per embryo in ICM and TE lineages.

**Supplemental figure 3: Mechanism of *Etv5* action on ICM cell fate decision (related to figure 3)**

**(A)** Quantification of inner cell mass (ICM) lineage composition in control and FGF treated wild-type, *Etv5^+/-^* and *Etv5^-/-^* embryos (related to Figure 3B). Double positive (DP; NANOG+GATA6+), primitive endoderm (PrE; NANOG-;GATA6+), epiblast (EPI; NANOG+;GATA6-light red, NANOG-;GATA6-dark red). Individual embryos shown and ordered by ascending cell number.

**Supplemental figure 4: Loss of *Etv4/5* causes a delay in the progression of pluripotency**

**(A)** Quantification of KLF4 and SOX2 levels in epiblast cells per embryo in an allelic *Etv4;Etv5* series of late-stage blastocysts (from Figure 4A).

**(B)** Quantification of KLF4 and SOX2 levels in epiblast cells per embryo in an allelic *Etv4;Etv5* series of late-stage blastocysts from (A) grouped by *Etv5* genotype.

**(C)** Confocal images of immunofluorescence staining of SOX2, OTX2 and F-actin in posti-implantation *Etv4;Etv5* day E5.5. embryos. Pr = Proximal, D = Distal, A = Anterior, P = Posterior. Scale bar 100μm

**(D-E)** Confocal images of immunostaining for OCT4, NANOG, KLF4 (D) and SOX2, OTX2, KLF4 (E) in wild-type and *Etv4^-/-^;Etv5^-/-^*mESC in naïve conditions (Serum/LIF) and differentiated to epi-like cells (EpiLC, FGF+Activin 48h). Scale bar 20μm

**Supplemental figure 5: Compound *Etv4/5* mutants have developmental delay and anterior visceral endoderm migration defects**

**(A)** Confocal images of an allelic series of *Etv4;Etv5* embryos at pre-streak stages, embryonic day (E) 6.25 immunostained for SOX2, and CER1. Anterior max intensity projection (right) and z-slice (left). Arrow heads show abnormal anterior/distal visceral endoderm migration and/or morphology. A = Anterior, P= Posterior. Pr = Proximal. D = Distal. Scale bar 100μm

**(B)** Confocal maximum intensity projection images of an allelic series of *Etv4;Etv5* embryos at late-bug gastrulation stages, embryonic day (E) 7.5 immunostained for SOX2, T and NANOG. Arrow head shows abnormal anterior/distal visceral endoderm migration and/or morphology. A = Anterior, P= Posterior. Pr = Proximal. D = Distal. Scale bar 100μm. Range of abnormal morphologies amongst *Etv4^+/-^;Etv5^-/-^*embryos (right).

**(C-D)** Observed and expected numbers from *Etv5^+/-^* **(C)** and *Etv4^+/-^;Etv5^+/-^* **(D)** heterozygous intercrosses at weaning, postnatal day (P) 21.

