## Supplemental Figures for "ETV4 and ETV5 Orchestrate FGF-Mediated Lineage Specification and Epiblast Maturation during Early Mouse Development"

### Simon et al Supplemental Figure 1

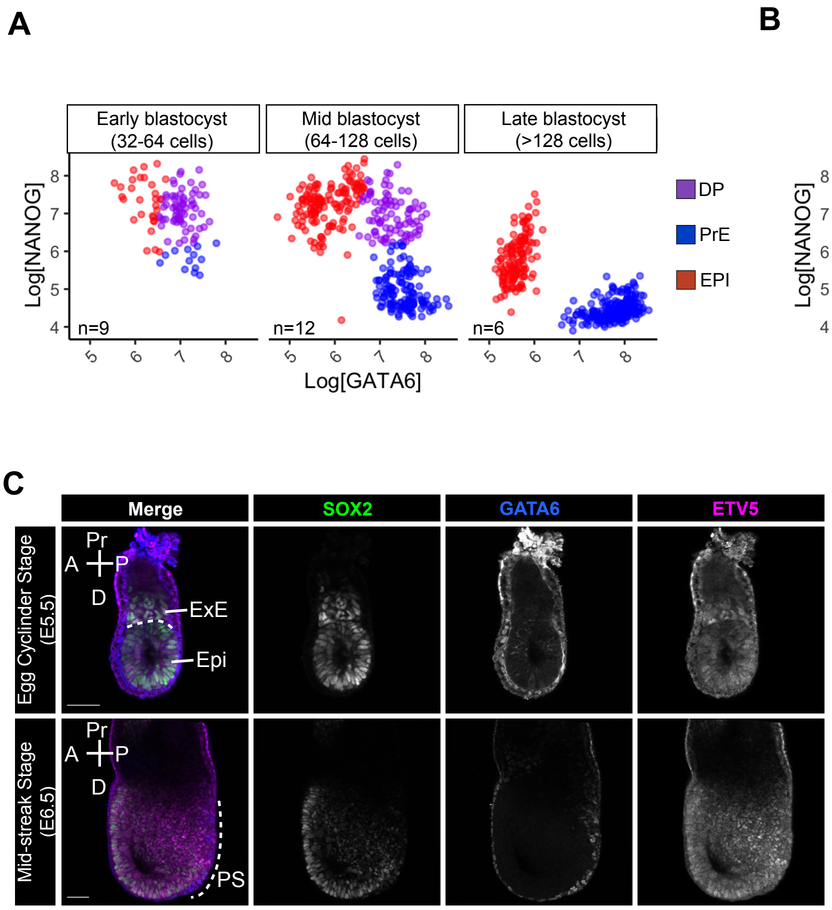

### Simon et al Supplemental Figure 2

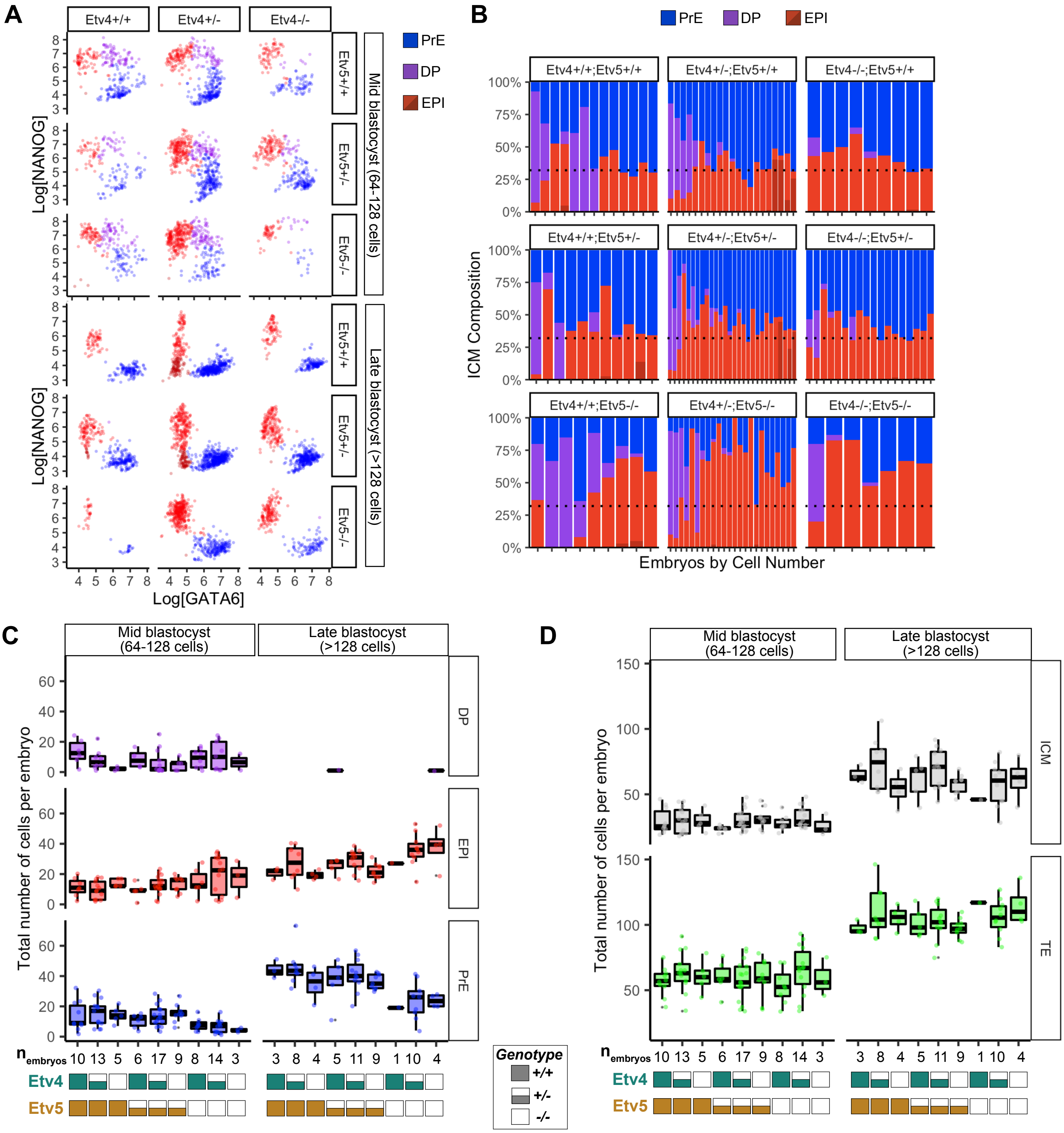

### Simon et al Supplemental Figure 3

**A**

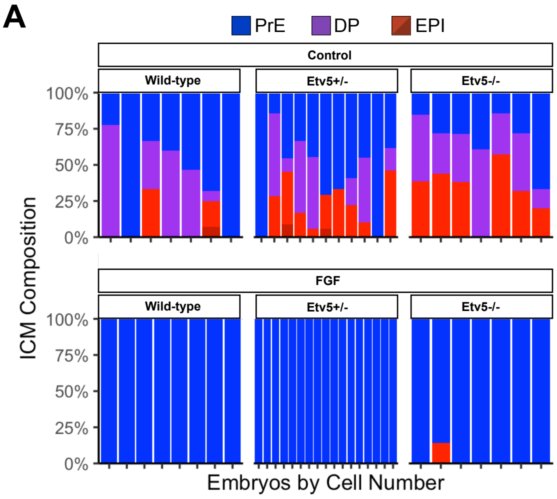

### Simon et al Supplemental Figure 4

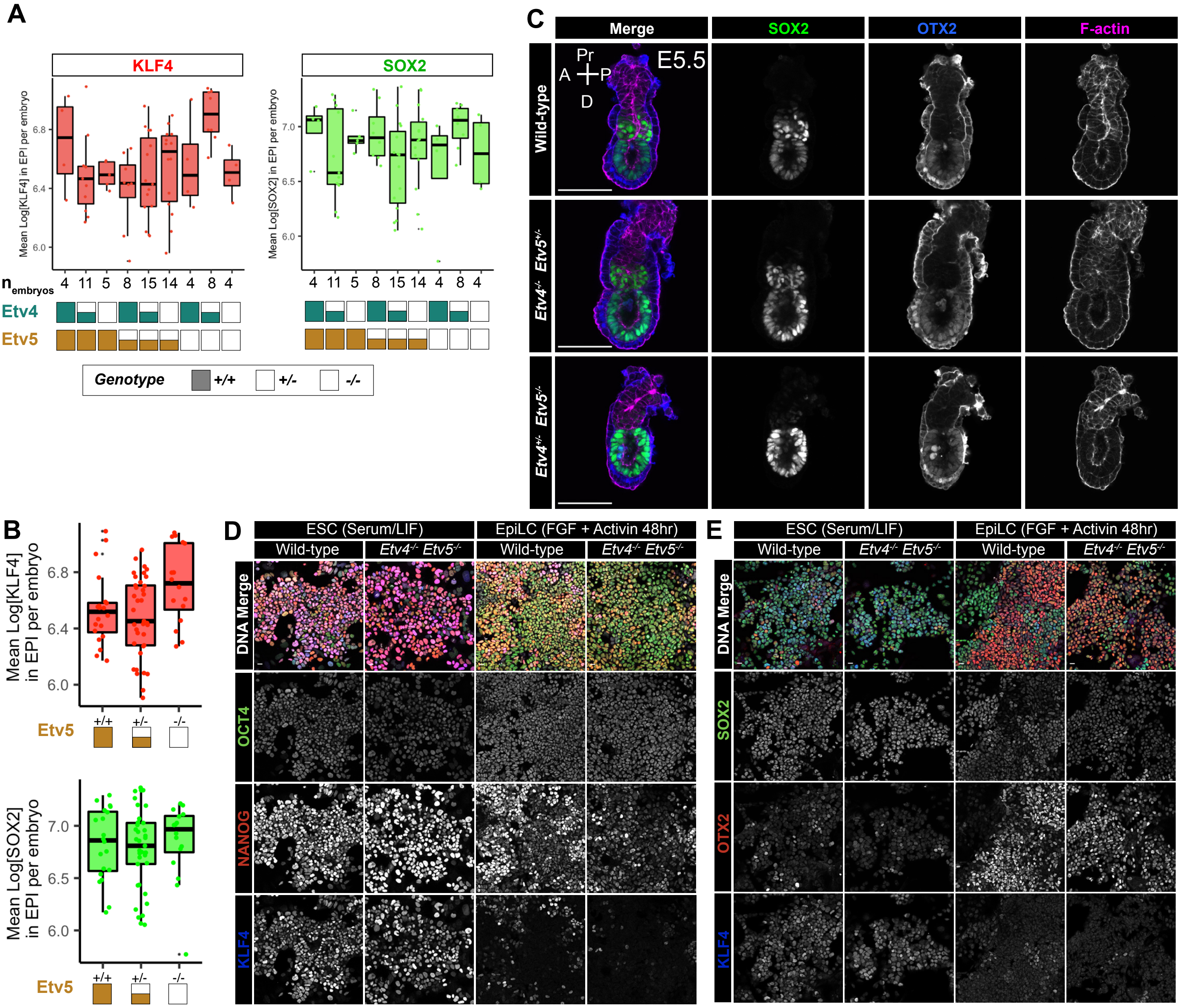

### Simon et al Supplemental Figure 5

**A**

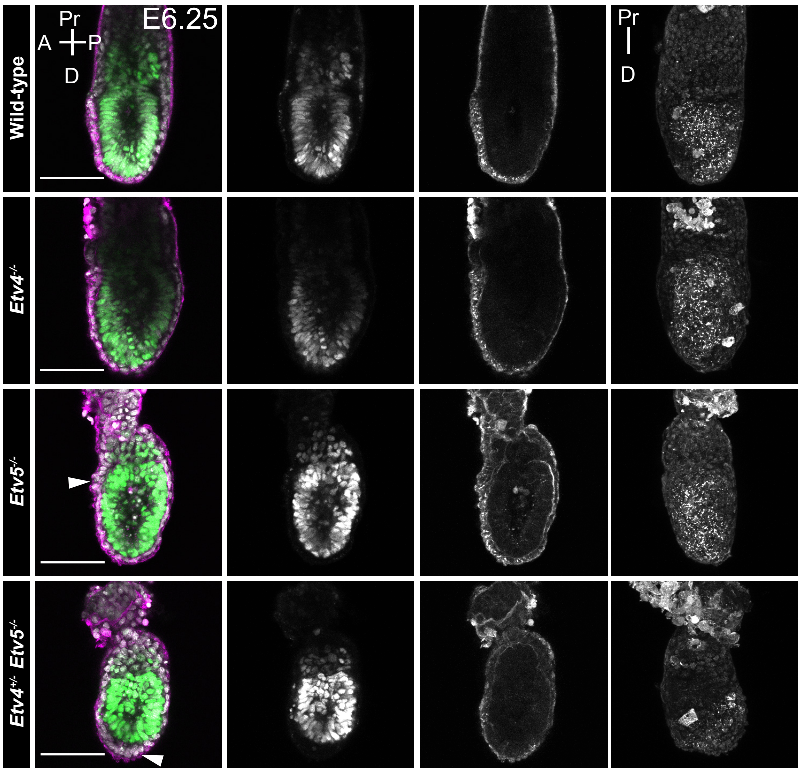


**B**

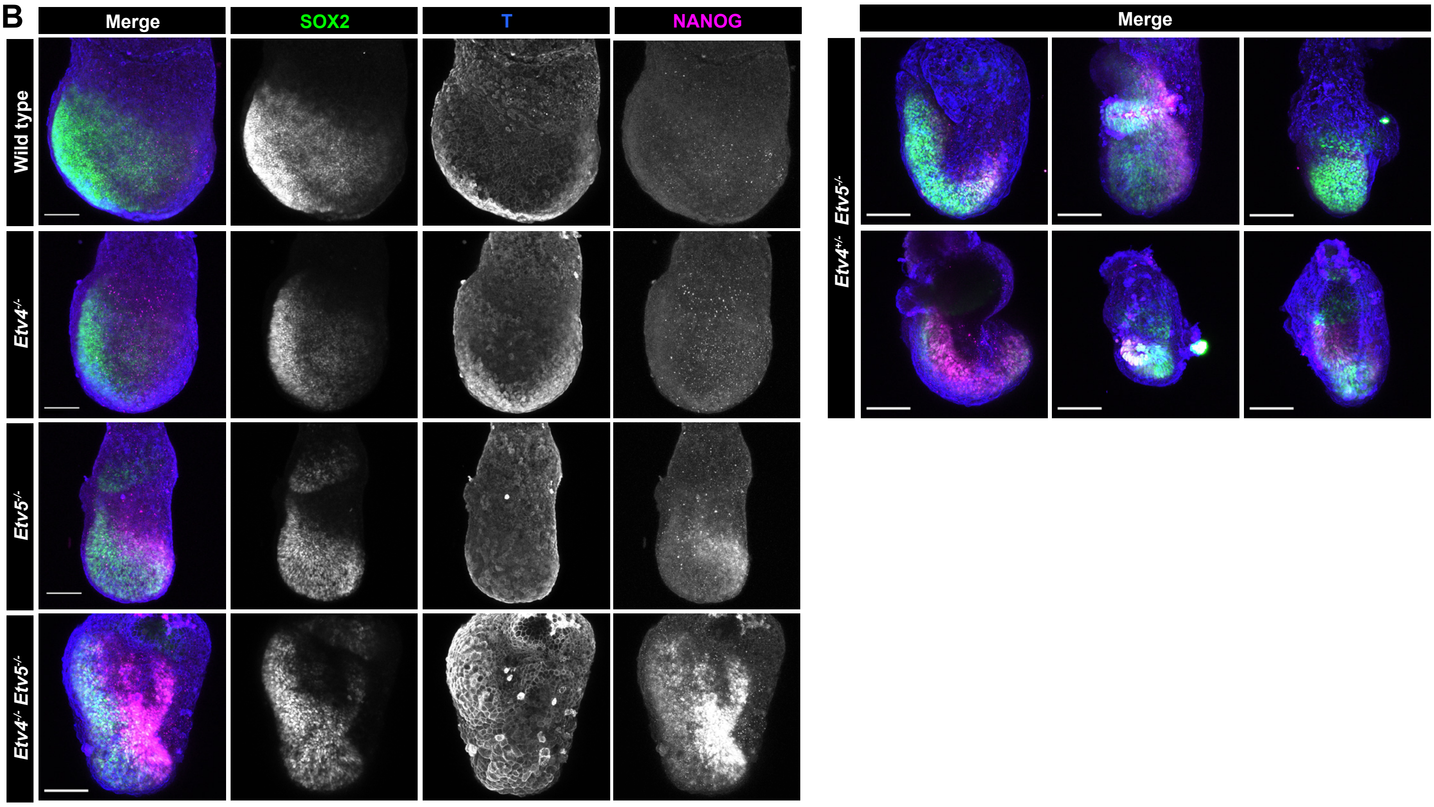


**C**

Observed *(Expected)* numbers: *Etv5*+/- intercross

| Stage | Genotype *Etv5* +/+ | +/- | -/- |
|---|---|---|---|
| **P21** | 19 *(16)* | 43 *(32)* | 1 *(16)* |

**D**

Observed *(Expected)* numbers: *Etv4*+/-;*Etv5*+/- intercross

Genotype: +/+ , +/- , -/- ; *Etv4*, *Etv5*

| Stage | | | | | | | | | |
|---|---|---|---|---|---|---|---|---|---|
| *Etv4* | +/+ | +/- | -/- | +/+ | +/- | -/- | +/+ | +/- | -/- |
| *Etv5* | +/+ | +/+ | +/+ | +/- | +/- | +/- | -/- | -/- | -/- |
| **P21** | 5 *(2)* | 4 *(2)* | 1 *(2)* | 4 *(4)* | 14 *(7)* | 0 *(4)* | 0 *(2)* | 0 *(4)* | 0 *(2)* |
